# GeneJepa: A Predictive World Model of the Transcriptome

**DOI:** 10.1101/2025.10.14.682378

**Authors:** Elon Litman, Tyler Myers, Vinayak Agarwal, Ekansh Mittal, Orion Li, Ashwin Gopinath, Timothy Kassis

## Abstract

We introduce GeneJepa, a self-supervised foundation model that learns a predictive world model of single-cell transcriptomes. Based on the Joint-Embedding Predictive Architecture, GeneJepa predicts latent representations of masked gene sets from visible context, a shift away from reconstructing noisy expression values and toward world-model style inference over cellular state. To realize this at scale, a Perceiver encoder handles variable gene sets at fixed cost, and a tokenizer jointly represents gene identity and continuous expression using Fourier features. Trained on the Tahoe-100M atlas, GeneJepa learns general representations that transfer across tissues and datasets. On downstream tasks, including drug response and perturbation prediction, it surpasses strong baselines and enables test-time scaling by progressively enlarging the cross-attention over the gene set, trading a small read cost for higher accuracy at inference. GeneJepa moves toward foundation models that reason over gene–gene relations, enabling applications in annotation, prediction, and in-silico discovery.

## 1 Introduction

Single-cell RNA sequencing (scRNA-seq) enables transcriptome-wide readouts for millions of individual cells, transforming our ability to chart cellular identity, state transitions, and responses to perturbation (Regev et al., 2017; Tabula Sapiens Consortium, 2022; Butler et al., 2018; Wolf et al., 2018; CZI Single-Cell Biology Program et al., 2023). Public and private atlases now span diverse tissues, species, and experimental conditions, while pooled CRISPR and small-molecule screens provide causal structure at unprecedented scale (Dixit et al., 2016; Adamson et al., 2016; Norman et al., 2019; Zhang et al., 2025). Yet the same properties that make these resources powerful—extreme dimensionality, sparsity from dropout, batch heterogeneity, and compositional confounding—continue to limit supervised pipelines and hand-engineered features (Lopez et al., 2018; Tran et al., 2020; Lähnemann et al., 2020).

Foundation models offer an alternative: learn transferable representations from unlabeled corpora, then adapt to downstream tasks with minimal supervision (Devlin et al., 2018; Brown et al., 2020; Radford et al., 2021; He et al., 2022). Several groups have proposed transcriptomic “foundation” models, most prominently scGPT and related Transformer-based variants (Cui et al., 2024; Yang et al., 2022a; Theodoris et al., 2023; Hao et al., 2024; Cui et al., 2022). These approaches typically tokenize genes (and often binned expression) into sequences, (ii) pretrain with masked token prediction or next-token objectives, and (iii) fine-tune for tasks such as annotation, perturbation effect prediction, and integration. While influential, token-level generative objectives inherit key mismatches with scRNA-seq: they emphasize exact reconstruction of noisy, zero-inflated counts; require arbitrary or ad hoc orderings over inherently set-structured inputs; and can overfit to batch- or lab-specific low-level statistics rather than abstract regulatory structure (Vallejos et al., 2017; Townes et al., 2019; Hafemeister and Satija, 2019; Luecken et al., 2022). In practice, masked reconstruction can yield strong in-domain metrics yet brittle transfer across labs, perturbation regimes, or quantification protocols (Luecken et al., 2022; Tran et al., 2020; Stuart et al., 2021).

**Our premise** is that models should not be judged on their ability to reproduce every read count, but on whether they capture the latent rules that govern co-expression programs and cellular state transitions. We therefore adopt the *Joint-Embedding Predictive Architecture* (JEPA) paradigm (LeCun, 2022; Assran et al., 2023a,b): rather than reconstructing raw inputs, the model learns to predict the *representation* of a masked target subset of genes from the *representation* of an observed context subset. This shift from signal reconstruction to *representation prediction* has three conceptual advantages over token generative modeling (e.g., scGPT): (1) it de-emphasizes incidental noise in count space and focuses learning pressure on abstract dependencies among genes and programs; (2) it removes the need for an arbitrary sequence order, naturally aligning with the set-like structure of expressed genes; and (3) it avoids contrastive negatives and likelihood-based calibration issues, yielding stable training at scale (LeCun, 2022; Bardes et al., 2021; Caron et al., 2021).

We instantiate this idea in GeneJepa, a predictive world model of the transcriptome. Architecturally, GeneJepa uses a Perceiver encoder to decouple input size from compute, allowing a fixed-cost latent array to attend over variable-length gene sets (Jaegle et al., 2021). Each gene–expression pair is mapped via a tokenizer that combines learnable gene embeddings with continuous expression encodings using Fourier features to preserve quantitative information without binning (Tancik et al., 2020; Rahimi and Recht, 2007). A student network predicts target embeddings for masked subsets given the context; a momentum (EMA) teacher produces targets, with variance–covariance regularization to prevent collapse and centering for stability (Tarvainen and Valpola, 2017; Bardes et al., 2021; Caron et al., 2021). Training on the large-scale Tahoe-100M perturbation corpus drives the model to capture causal regularities that generalize across tissues, technologies, and interventions (Zhang et al., 2025).

### Relation to prior work

Compared with scGPT-style masked token models (Cui et al., 2024), GeneJepa (i) predicts in embedding space rather than reconstructing noisy counts, (ii) treats inputs as unordered sets queried by a latent bottleneck, avoiding positional heuristics, (iii) removes likelihood calibration and vocabulary design (no expression binning), and (iv) scales with stable, non-contrastive objectives (LeCun, 2022; Bardes et al., 2021). Unlike scGPT and other Transformer pretraining on cell corpora (Cui et al., 2024; Rosen et al., 2023), GeneJepa‘s target-in-embedding objective also explicitly optimizes for *predictive sufficiency* of context over targets, encouraging the model to encode regulatory programs and conditional independencies rather than memorizing token frequencies or batch artifacts. Finally, unlike contrastive pipelines that rely on handcrafted augmentations and negatives (Chen et al., 2020; Yang et al., 2022b), GeneJepa learns from natural co-occurrence structure in real cells, side-stepping augmentation design pitfalls in genomics.

### Contributions

1. **Paradigm**. We introduce JEPA-based representation prediction for scRNA-seq, arguing and demonstrating that abstract predictive objectives better align with noisy, set-structured transcriptomes than token reconstruction or contrastive learning (LeCun, 2022; Assran et al., 2023b).
2. **Architecture & Practical Innovations**. We design a scalable, set-aware encoder with Perceiver latents and a Fourier-feature tokenizer for continuous expression, paired with an EMA teacher and variance–covariance regularization for stable training (Jaegle et al., 2021; Tancik et al., 2020; Tarvainen and Valpola, 2017; Bardes et al., 2021; Caron et al., 2021).
3. **Evidence**. Trained on Tahoe-100M, GeneJepa yields embeddings that transfer across datasets, labs, and assays; enable linear probes for annotation; and support perturbation reasoning with strong out-of-domain robustness, surpassing generative token baselines including scGPT in accuracy and calibration on held-out regimes (Zhang et al., 2025; Luecken et al., 2022; Tran et al., 2020; Cui et al., 2024).

GeneJepa moves beyond reconstructing expression counts and instead learns to predict the abstract regulatory state of held-out genes. This change in objective produces models that capture the fundamental rules of cellular systems and generalize across the diverse landscape of single-cell biology (LeCun, 2022; Norman et al., 2019; Regev et al., 2017).

## 2 The GeneJepa Architecture

GeneJepa is a way of doing self-supervision for single-cell transcriptomics that follows the Joint-Embedding Predictive Architecture. The model has three parts: a student encoder, a momentum based teacher encoder, and a predictor. The goal is straightforward:

From a subset of genes in a cell, the student produces a context representation, and the predictor uses this context to forecast the representation of a different subset of genes. The teacher provides the target representation that the predictor aims to match.

The procedure, shown in Figure 1, starts from one cell’s expression profile. We treat the profile as a set of (gene, value) pairs and split it at random into two disjoint sets: a *context block* and a single *target block*. The context block is passed through the student encoder *f*_***θ***_ to produce a context representation ***z***_ctx_. The predictor *p*_***ϕ***_ then takes this context representation as its sole input and outputs a predicted target representation, 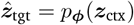.

**Figure 1.**
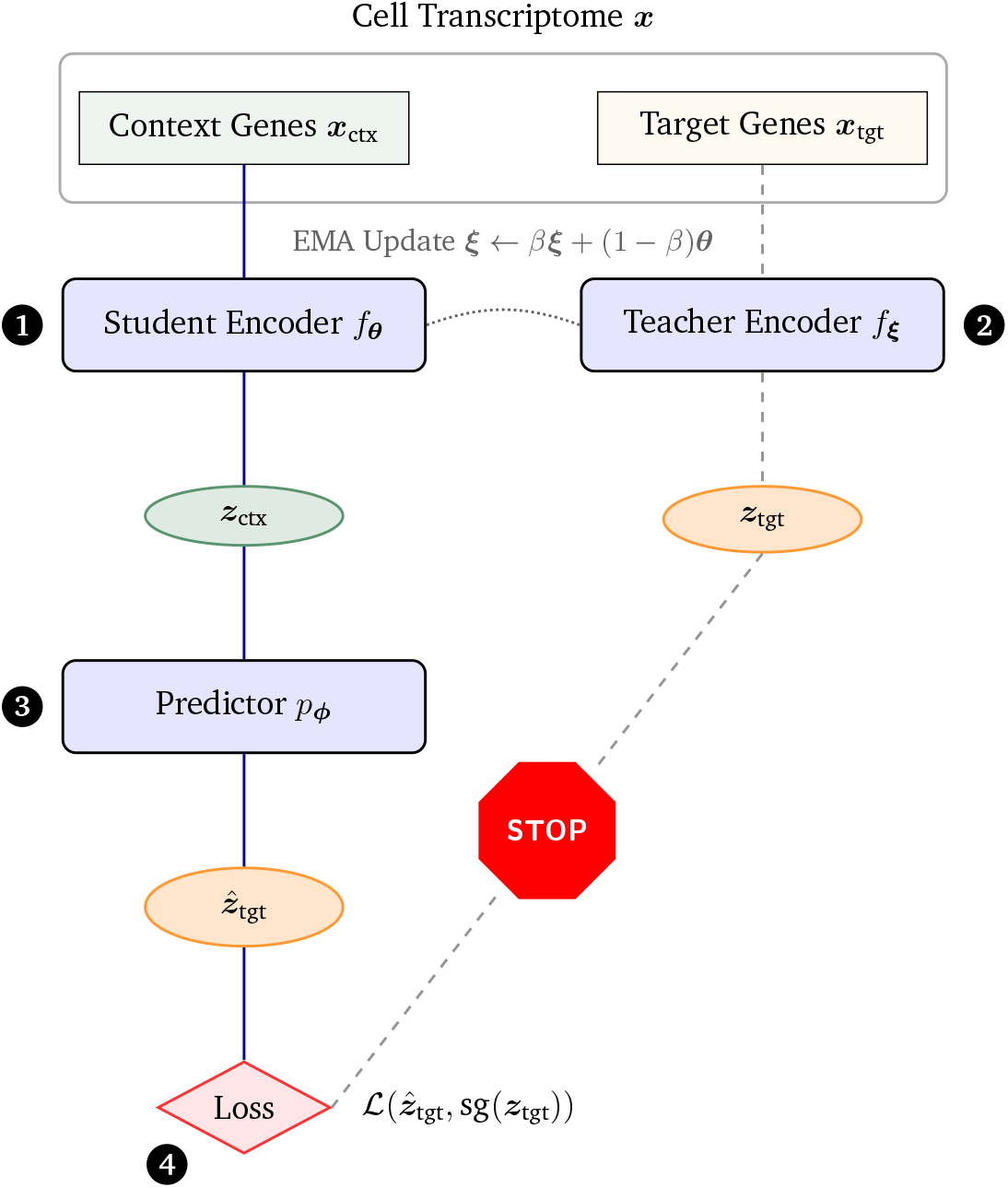
The GeneJepa Architecture. A single cell’s transcriptome, ***x***, is partitioned into a contex block, ***x***_ctx_, and a target block, ***x***_tgt_. ❶ The context genes are tokenized and processed by the **student encoder** (*f*_***θ***_) to produce a context representation ***z***_ctx_. ❷ The target block is processed by the momentum **teacher encoder** (*f*_***ξ***_) to produce the target representation ***z***_tgt_. The teacher’s weights (***ξ***) are an Exponential Moving Average (EMA) of the student’s weights (***θ***). ❸ A **predictor** network (*p*_***ϕ***_) takes the context representation ***z***_ctx_ and predicts the target representation, yielding 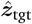. ❹ The **loss** is computed to minimize the distance between the prediction 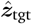 and the target ***z***_tgt_. The gradient is stopped from flowing back into the teacher network (indicated by the dashed line, the `sg`(·) notation, and the explicit STOP sign), ensuring the student learns to predict a stable, informative target.

In parallel, the genes in the target block are processed by the teacher encoder *f*_***ξ***_ to produce the groundtruth target representation ***z***_tgt_ = *f*_***ξ***_(target_block). The teacher shares the student’s architecture, but its weights ***ξ*** are an exponential moving average (EMA) of the student’s weights ***θ***. We stop the gradient from flowing into the teacher, making it a stable, slowly evolving target for the student to predict.

Training minimizes the discrepancy between the prediction 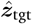 and the target ***z***_tgt_ in a shared embedding space. By predicting a large, held-out portion of the transcriptome from the context, the student is forced to learn a holistic representation of cellular state. This encourages the model to infer the latent dependencies between genes, which is the central predictive task that defines GeneJepa.

### 2.1 Tokenization

A challenge in applying deep learning to scRNA-seq is engineering an effective input representation. A cell’s state is defined not only by which genes are expressed but also by their expression levels. Discretizing expression values into bins, a common practice in prior work, discards fine-grained quantitative information. To address this, we designed a custom tokenizer that jointly models gene identity and continuous expression values.

Given a gene with vocabulary index *i* and a log-normalized expression value *v*, our tokenizer produces a token embedding ***t*** *∈* ℝ^*d*^. This is achieved in three steps. First, the gene’s identity is mapped to an embedding vector 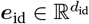 via a standard learnable embedding layer: ***e***_id_ = Embedding(*i*).

Second, the continuous expression value *v* is encoded into a high-dimensional vector 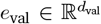 using Fourier features.

#### Definition 2.1

Numerical Fourier Tokenization

This technique projects a scalar value onto a set of sinusoidal basis functions, allowing the network to easily reason about continuous inputs. We define a set of *N*_*f*_ frequencies, 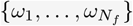, logarithmically spaced between a minimum and maximum frequency. The value embedding is then computed as:

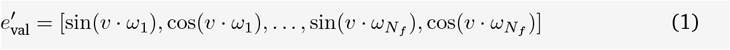

This raw Fourier embedding 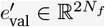 is then processed by a small multi-layer perceptron(MLP) to produce the final value embedding *e*_val_ of the desired dimension *d*_val_.

Third, the identity and value embeddings are concatenated, [***e***_id_, ***e***_val_], and passed through a final projection layer with Layer Normalization and a GELU activation to produce the final token embedding ***t*** ∈ ℝ^*d*^, where *d* = *d*_id_ + *d*_val_. This tokenizer creates a rich, continuous representation for every expressed gene in a cell.

### 2.2 Perceiver Encoder

The student and teacher share the same backbone, a Perceiver encoder. We use the Perceiver because single-cell profiles contain a variable number of expressed genes, and the model must process inputs whose length can change from cell to cell without a blow-up in compute or memory. The Perceiver achieves this by separating “how much you read” from “how much you think”: it reads the variable-length input with attention, then thinks over a fixed-size latent state.

#### Cross-attention

Let 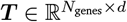 be the tokenized genes for one cell and 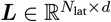 be a learned latent array of fixed size. The encoder first lets the latents query the input tokens through a single cross-attention layer, which compresses the variable-length set of genes into the fixed latent array. We write this update as

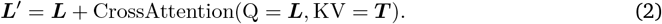

To cope with very large *N*_genes_ without materializing a full attention matrix, we compute this operation in chunks over the keys and values using a numerically stable online softmax. The core softmax arithmetic is kept in `float32` to avoid underflow or overflow during mixed-precision training; see A.2 and (Milakov and Gimelshein, 2018; Dao et al., 2022; Rabe and Staats, 2021).

#### Latent processing

The updated latents ***L***^*′*^ are then passed through a deep stack of Transformer blocks that apply self-attention only among the latent vectors. Because *N*_lat_ is fixed, the cost of this “thinking” stage does not grow with the number of expressed genes, which lets us use a deeper stack to capture richer interactions. We apply gradient checkpointing across these blocks to further control memory use during training.

#### Global readout

After the latent stack, we aggregate the final latent vectors by mean pooling to obtain a single representation ***z*** ∈ ℝ^*d*^ for the cell or gene subset. This vector is the encoder’s output and is used as the context embedding ***z***_ctx_ or as the target embedding ***z***_tgt_, depending on which genes were provided to the encoder.

### 2.3 Masking Strategy and Predictor

The predictive task in GeneJepa is defined by its masking strategy, which partitions a cell’s transcriptome into a context set and a target set. For each cell, we first randomly permute its set of expressed genes to remove any potential for the model to learn spurious order-based dependencies. We then designate a fraction of these genes (around 45%) as the target block, with the remaining genes forming the context block. To ensure that the context is sufficiently informative and the target is non-trivial, we enforce minimum size constraints on both sets. In our experiments, we found that generating a single, large target block per cell provided a strong and stable training signal.

The predictor is a feed-forward network whose goal is to predict the teacher’s representation of the target block, ***z***_tgt_, using only the student’s representation of the context block, ***z***_ctx_. Because each context is paired with a single target, the prediction task is unambiguous, and no additional positional or content-based information about the target is required. The predictor, a deep Multi-Layer Perceptron (MLP) with GELU activations and Layer Normalization, therefore takes only the context representation ***z***_ctx_ as input.

The predictor receives no explicit information, such as gene identities, about the target block it is tasked with predicting. This design choice prevents the model from learning trivial shortcuts or cheating via information leakage. It forces the student encoder to distill a holistic representation of the cell’s state into the context embedding, ***z***_ctx_, from which the latent state of the held-out genes can be inferred. This demanding objective encourages the development of representations that capture the fundamental rules of gene co-regulation. The predictor’s output is the predicted target representation:

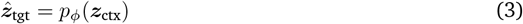

### 2.4 Teacher Network

The teacher network provides the ground truth targets for the predictor. It is structurally identical to the student encoder but is not updated via backpropagation. Instead, its weights, ***ξ***, are an exponential moving average (EMA) of the student’s weights, ***θ***. At each training step *k*, the update rule is:

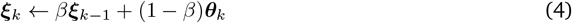

where *β* is the EMA decay rate. This momentum mechanism ensures that the teacher evolves more slowly than the student, providing a stable and consistent set of targets. This prevents the trivial solution where the student and teacher chase each other in the embedding space and ultimately collapse. The decay rate *β* is not fixed but is scheduled to increase over the course of training, starting from a lower value (e.g., 0.996) and annealing towards a value very close to 1 (e.g., 0.9995) using a cosine schedule. This allows the teacher to track the student more closely in the early stages of training and become more stable as the student converges.

## 3 Objective Function & Training

The training of GeneJepa is guided by a carefully constructed objective function designed to encourage accurate prediction while preventing various forms of representational collapse. The overall training procedure is implemented on a massive scale, leveraging efficient data loading and distributed training techniques.

### 3.1 Loss Formulation

The training of GeneJepa is guided by a composite objective function designed to enforce accurate prediction in the latent space while simultaneously preserving the informational richness of the learned representations. The total loss is a weighted sum of a primary predictive loss and a regularization term that explicitly prevents representational collapse.

#### Predictive Loss

The central learning signal is derived from the dissimilarity between the predictor’s output, 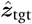, and the teacher’s target representation, ***z***_tgt_. We use a cosine similarity loss, which measures the alignment between the two vectors, making the objective invariant to their magnitudes. This encourages the model to capture the correct semantic direction in the embedding space without being overly constrained by vector length. For a given prediction, the loss is formulated as:

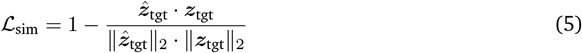

Crucially, a stop-gradient is applied to the teacher’s output ***z***_tgt_. This ensures that gradients flow only into the student and predictor networks, preventing the trivial solution where the teacher network is modified to simply match the student’s (potentially incorrect) predictions.

#### Collapse Prevention with VICReg

A significant challenge in self-supervised models that do not rely on explicit negative samples, such as Jepa, is the risk of *representational collapse*. This occurs when the network learns a trivial solution by outputting a constant vector regardless of the input. To counteract this, we incorporate the Variance-Invariance-Covariance Regularization (VICReg) loss (Bardes et al., 2021) directly on the batch of predictions, 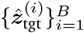. This regularizer is composed of two key terms that work in concert to maintain the quality of the embedding space.

The **variance** term addresses feature collapse, where embeddings collapse to a single point. It encourages the standard deviation of the embeddings along each dimension to approach a target value of 1, thereby ensuring that the model utilizes the embedding space. The loss for the *j*-th dimension is:

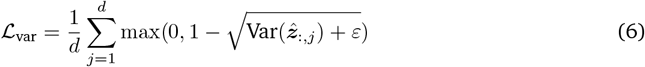

where 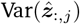 is the variance of the *j*-th feature across the batch and *ε* is a small constant for numerical stability.

The **covariance** term addresses dimensional collapse, where the learned features are highly correlated and thus informationally redundant. This term penalizes the off-diagonal elements of the covariance matrix of the embeddings, effectively decorrelating the feature dimensions and encouraging the model to spread information across the entire representation vector.

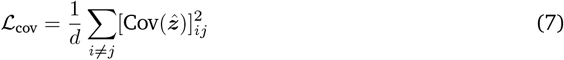

By combining these terms, VICReg ensures that the predictions lie on a well-behaved manifold, preventing both feature-level and dimensional collapse without requiring a momentum-based centering mechanism. The final objective function is a linearly weighted sum of these three components, balancing the predictive accuracy with the structural integrity of the representation space:

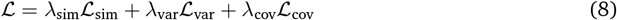

where *λ*_sim_, *λ*_var_, and *λ*_cov_ are scalar hyperparameters that control the relative importance of each term.

### 3.2 Large-Scale Implementation Details

The scale of the Tahoe-100M dataset poses significant data handling and computational hurdles. Our implementation addresses these through on-disk data streaming and mixed-precision distributed training.

#### Dataset and Preprocessing

We train GeneJepa on the Tahoe-100M dataset (Zhang et al., 2025), which contains over 100 million single-cell profiles from approximately 1,000 cancer cell lines subjected to over 3,000 small-molecule compound perturbations. This dataset provides immense diversity in cellular states and tissue type. To ensure stable and consistent normalization, we precompute global statistics (mean and standard deviation) from a large subset of the training data. All expression counts are transformed with log(1 + *x*) and then standardized using this global Z-score normalization. This is a critical step that avoids the batch-to-batch statistical noise inherent in using batch normalization for input data.

#### Efficient Data Loading

We use the Hugging Face datasets library to stream the `dataset` directly from disk, avoiding the need to load the entire dataset into memory. Our custom DataModule, built using PyTorch Lightning, handles the DDP-safe sharding of data files across multiple GPUs and workers. For training, the data stream is shuffled using a large buffer to ensure randomness, while for validation, shuffling is disabled for reproducibility.

#### Training Configuration

The model is trained using the PyTorch Lightning framework, which automates distributed training, mixed-precision, and logging. We use the AdamW optimizer with a weight decay of 2 × 10^−4^ and a cosine learning rate schedule with a linear warmup phase. Onedimensional parameters (such as norm scale and biases) and embedding tables are excluded from weight decay. Training is performed on a cluster of 4 NVIDIA H100 80GB GPUs using `bfloat16` mixed-precision to accelerate computation and reduce memory usage. The Distributed Data Parallel (DDP) strategy is used to synchronize gradients across devices, with the `find_unused_parameters` flag set to True to accommodate the model’s architecture where not all parameters may be used in every forward pass due to dynamic masking. For a complete list of hyperparameters and model configurations, please see Appendix A.

## 4 Results

To evaluate the representations learned by GeneJepa, we conducted a series of experiments on diverse, unseen downstream tasks. We benchmarked our model against leading foundation models for single-cell genomics, including the generative model scGPT (Cui et al., 2024) and Universal Cell Embeddings (UCE) (Rosen et al., 2023). All backbone encoders (GeneJepa, scGPT, UCE) were used as frozen feature extractors with no fine-tuning; for ridge readouts we tuned *α* on the same log-spaced grid via five-fold CV using the identical protocol across backbones.

### 4.1 Representations Capture Biological Identity Across Tissue

A key measure of a foundation model’s success is its ability to organize cells in its embedding space according to their fundamental biological identity, without having been trained on explicit labels. We tested this capability on two complex atlases.

#### 4.1.1 Peripheral Blood Mononuclear Cells 3k

We first assess whether GeneJepa‘s representation separates the basic immune cell types we expect to see in blood. For this we use the standard Peripheral Blood Mononuclear Cells benchmark with about three thousand cells. It is small, clean, and contains cells such as B cells, CD4 and CD8 T cells, NK cells, monocytes, dendritic cells, and platelets. Because it is uncomplicated, it is a good place to see whether a model learns the right geometry without any task-specific training.

We embed all cells with frozen encoders. Labels are created once by clustering and then mapping clusters to canonical immune types with marker genes. These labels are not used to shape the embeddings. They serve only to measure what the geometry already contains. For visualization we fit UMAP on the test embeddings only, separately for each method, so that the plots reflect the geometry induced by the encoder rather than the downstream classifier. For quantitative evaluation we train two very simple readers on the training split of the embeddings, a logistic regression classifier and a cosine k-nearest neighbors classifier, and we report Macro F1 and Accuracy on the held out split. Splits, preprocessing, and readouts are matched across methods.

Figure 2 shows visualizations and evaluations on the embeddings. In Figure 2A the GeneJepa test embeddings form compact and well separated groups that correspond to B cells, CD4 and CD8 T cells, natural killer cells, monocyte subsets, dendritic cells, and platelets. In Figure 2B the scGPT test embeddings appear as scattered strands with interleaving of types, which suggests a weaker organization of identity. Figure 2C shows the linear probe on GeneJepa reaches a Macro F1 of 0.69 with an Accuracy of 0.68 on the same fixed split. The k-nearest neighbors reader shows the same trend with a Macro F1 of 0.38 and an Accuracy of 0.52. The scGPT readers are lower in both cases with 0.23 and 0.20 for the linear probe and 0.25 and 0.37 for k-nearest neighbors. Figure 2D reports per class F1 and shows that the advantage for GeneJepa is not confined to a single abundant type. Gains are visible for common lymphoid populations and also for rarer platelets and dendritic cells.

**Figure 2.**
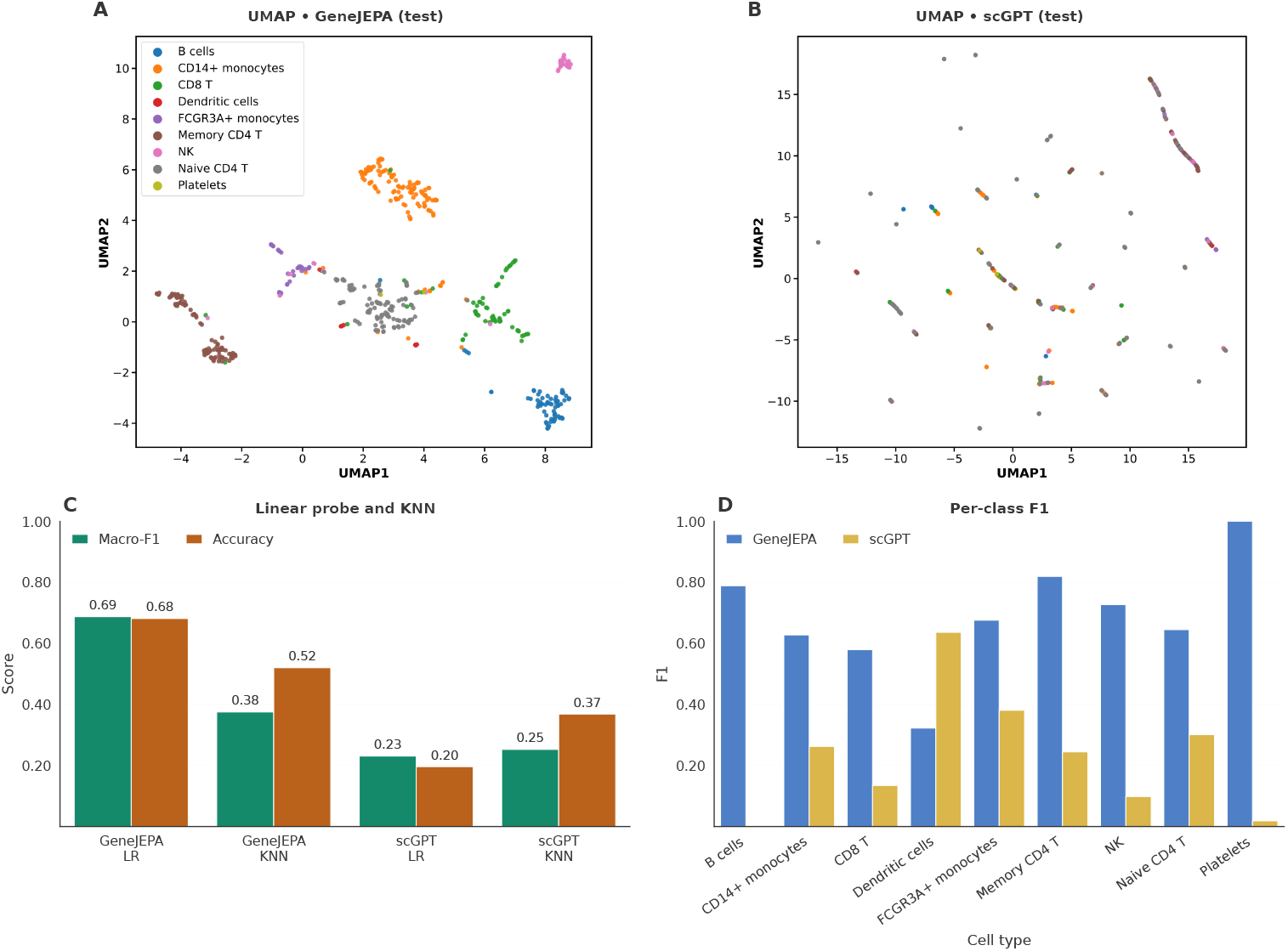
GeneJepa learns a superior manifold of cell identity on PBMC3k. **(A)** UMAP of GeneJepa test embeddings shows compact, well-separated immune groups. **(B)** UMAP of scGPT test embeddings shows fragmented, interleaved structure. **(C)** Linear-probe and kNN results (Macro-F1, Accuracy) on the same split favor GeneJepa. **(D)** Per-class F1 highlights consistent gains, including on rarer types.

#### 4.1.2 Human Lung Cell Atlas

We assessed representation quality on the Human Lung Cell Atlas (HLCA) (Sikkema et al., 2023), which comprises healthy lung cells spanning dozens of fine-grained epithelial, endothelial, stromal, and immune types. All cells were embedded with the frozen encoder. To quantify how well the embedding geometry reflects biological identity, we performed two complementary evaluations.

First, we fit a linear probe: a multiclass logistic regression with *ℓ*_2_ regularization trained on the embeddings to predict the HLCA annotations. The train/validation/test split was stratified by cell type (80/10/10), and all hyperparameters were selected on the training set using five-fold cross-validation. We report Macro F1 (to weight rare types fairly) and overall accuracy on the held-out test set. Second, we examined unsupervised structure by running *k*-means on the test embeddings with *k* set to the number of annotated types present in the split. We visualized the resulting manifold with UMAP fit *only* on test embeddings, and we summarize alignment between clusters and annotations via the confusion matrix as well as per-class F1 scores. Figure 3 shows linear separability, unsupervised concordance with biology, and error structure across abundant and rare populations.

**Figure 3.**
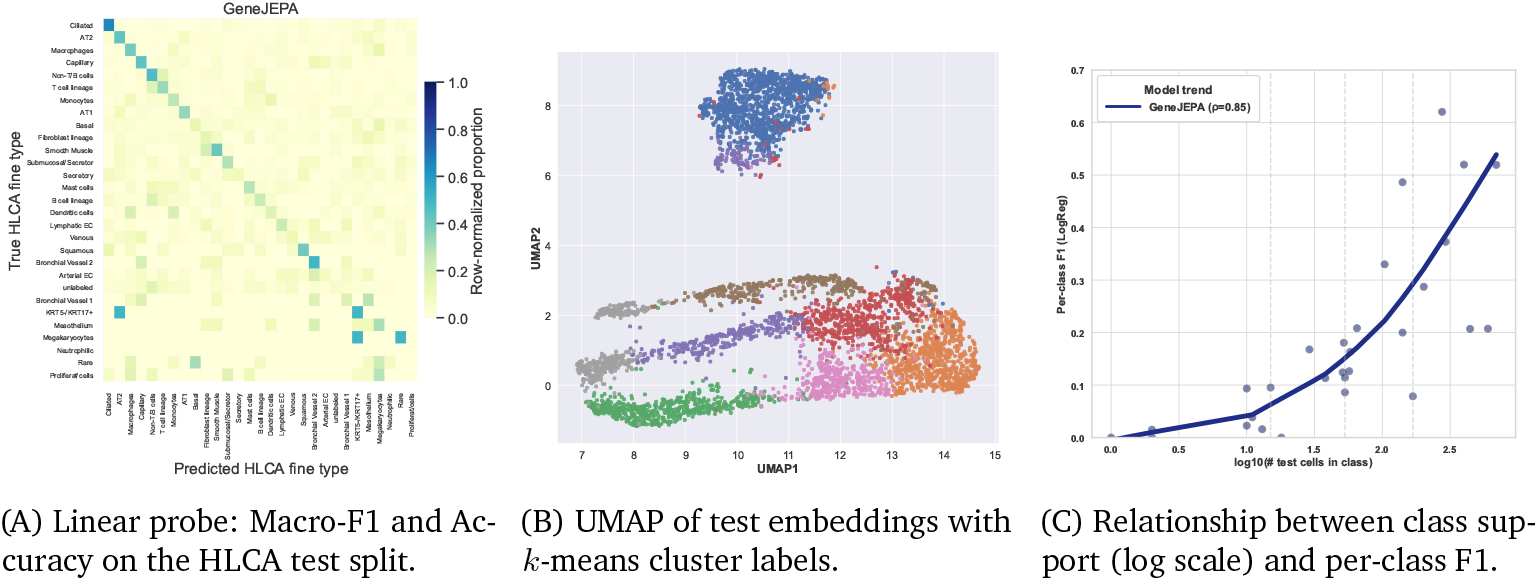
GeneJepa learns a robust manifold of cell identity on the Human Lung Cell Atlas. **(A)** Confusion matrix from a linear probe trained on frozen embeddings and evaluated on the held-out test set. The strong diagonal indicates high classification accuracy across most cell types. **(B)** UMAP visualization of the test set embeddings, with colors corresponding to unsupervised k-means clusters. The plot reveals a well-structured manifold where cell populations form distinct, well-separated groups. **(C)** Per-class F1 score as a function of class support (number of cells, log10 scale). Performance strongly correlates with class size (Spearman’s *ρ* = 0.85), demonstrating that the model learns more robust representations for more abundant cell types.

### 4.2 Drug Response

We evaluate single-cell dose-response prediction as a regression problem. Given a cell’s transcriptomic profile, the goal is to predict the (log-transformed) compound dose associated with that cell.

#### Data and splits

We use the sci-Plex dataset (v3), following the standard preprocessing used across our benchmarks. For regression, we aggregate cells into pseudobulk profiles by available context keys (cell line, compound, time), which improves stability without relying on labels from the test set. We randomly partition the aggregated examples into train and test splits (80/20), ensuring all models see the *same* split.

#### Models

We compare GeneJepa to representative single-cell foundation backbones, including scGPT and UCE. Each model produces a fixed-dimensional embedding per example; a linear ridge regressor (RidgeCV with a log-spaced grid) is then fit on the training embeddings and evaluated on the held-out test embeddings.

#### Metrics

We emphasize error and robustness: root mean squared error (RMSE), mean absolute error (MAE), median absolute error (MedAE), and interquartile-normalized RMSE (NRMSE-IQR), all computed on the test set. To contextualize absolute error, we report a *baseline-normalized* RMSE (rRMSE) relative to a global-median predictor (values *<* 1 indicate improvement over the naive baseline). We also summarize *cross-context robustness* via the median per-context MAE and its IQR (lower is better), where contexts are the (cell line, compound, time) groups. Finally, we report the magnitude of mean bias (|mean(*y* − *ŷ*)|). Embeddings are computed once per model using identical inputs (same counts matrix, gene budget, and split). RidgeCV searches *α* over a log grid and uses five-fold cross-validation on the training set only. All metrics are computed on the frozen test set.

#### Results

Across error-focused and robustness-focused measures, GeneJepa is best: it achieves the lowest RMSE, MAE, MedAE, and NRMSE-IQR; it is the *only* model with rRMSE below the global-median baseline; and it exhibits both the lowest median per-context MAE and the tightest per-context error spread, with the smallest absolute bias. Figure 4 summarizes these comparisons.

**Figure 4.**
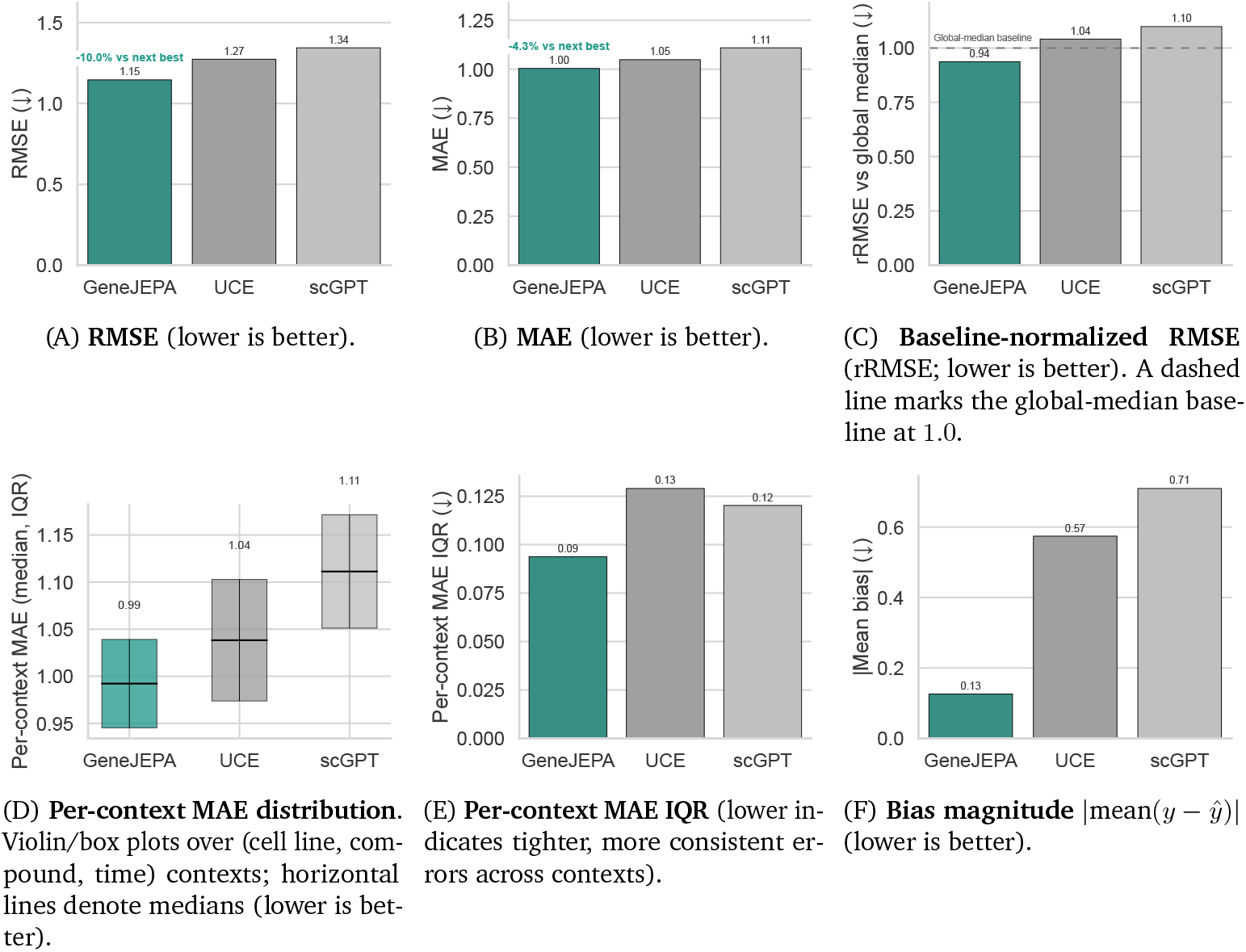
Drug-response regression on sci-Plex. Each model provides fixed embeddings; a ridge regressor is trained on the shared training split and evaluated on the held-out test set. **(A–B)** *Overall error:* GeneJepa attains the lowest RMSE and MAE among all baselines. **(C)** *Baseline-normalized error:* GeneJepa is the only model with rRMSE *<* 1, indicating improvement over a naive global-median predictor (dashed line). **(D–E)** *Cross-context robustness:* GeneJepa shows the lowest median per-context MAE and the tightest per-context error spread (IQR). **(F)** *Calibration of central tendency:* GeneJepa exhibits the smallest absolute mean bias. Bars display point estimates; where shown, thin lines denote summary statistics (median or reference line). All models use identical data preprocessing, gene budget, and train/test split; ridge hyperparameters are tuned only on the training set. GeneJepa delivers consistently lower error and tighter cross-context robustness than baseline alternatives, achieving the only sub-baseline normalized RMSE (0.94 ×), the lowest per-group median error (0.99), and the smallest bias.

### 4.3 Perturbation Prediction

Human biology involves the coordinated activity of approximately 37 trillion cells across tissues and organs. Single-cell technologies now resolve DNA, RNA, and protein variation at cellular resolution, yet mapping how small molecule perturbations causally alter cell state remains costly and limited in coverage. If we could predict transcriptional responses in new primary cell types from data, we could triage candidates and focus expensive assays where they matter most. Prior work on drug response modeling has largely centered on autoencoder families such as Dr. VAE, scGEN, and ChemCPA. These systems are commonly trained on the Connectivity Map, which contains more than 1.3 million perturbations but measures only 978 landmark genes and is dominated by immortalized cancer lines, limiting biological generalization. The Open Problems NeurIPS 2023 task addresses these issues by releasing a new single-cell perturbational dataset in primary human PBMCs with 144 compounds, three donors, treatment at 24 hours, and multiomic baseline measurements designed to inform cell type specific priors. The benchmark asks for the predicted post treatment gene expression profile for each compound and cell type.

#### Benchmark protocol

We adopt the official file contracts and metrics of the Open Problems benchmark. Training and test matrices are stored as `AnnData` objects with layers containing differential expression statistics. The prediction target for the benchmark is a directional significance score, *S*, defined as:

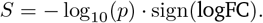

These scores are subsequently clipped to the interval [−4, 4]. We ensure that both the train and test matrices contain the clipped target layer. The gene order is fixed by `var_names` and the test row order is defined by the provided `id_map`. We evaluated with the five official row-wise metrics: cosine similarity, Pearson correlation, Spearman correlation, mean absolute error, and root mean squared error. In all experiments reported here, the test set contains 255 profiles and 18,211 genes.

#### GeneJepa-based probe

We benchmark a single pass probe built on the frozen GeneJepa encoder trained on single-cell RNA data. For each cell type, we compute a cell embedding by feeding a pseudo-expression vector into the teacher encoder of GeneJepa. The pseudo-expression vector is constructed from the training differential expression matrix by averaging the positive part of the log-fold change across rows that match the cell type. For each compound, we compute a chemical fingerprint using the Morgan procedure with radius two and 2,048 bits. We concatenate the cell embedding, the fingerprint, and the logarithm of the dose in micromolar to form a feature vector. A multi-output ridge regressor is trained on the training set to map features to the clipped signed log ten signal for all genes. The ridge parameter is selected on a held-out split that respects compound and cell type grouping. After fitting, we apply an affine calibration per gene on the training set to correct magnitude bias and then clip predictions to the benchmark support. For the results below, we report the version of the probe prior to calibration, which we refer to as GeneJepa, since it provides a clear picture of the directional signal provided by GeneJepa embeddings without magnitude adjustment.

#### Results

On the PBMC benchmark, the GeneJepa probe achieves a mean row-wise cosine of 0.3698, a mean row-wise Pearson of 0.3509, and a mean row wise Spearman of 0.3431. These values exceed a set of widely used controls and practical, widely-used baselines. The gains over the mean baselines indicate that GeneJepa embeddings capture biologically meaningful directions of change that generalize across compounds and cell types. The improvement over a Transformer ensemble on these directionality metrics is notable given the minimal probe. The results suggest that joint embedding predictive pretraining on scRNA-seq data provides transferable structure for drug response modeling in primary immune cells.

### 4.4 Test-Time Scaling in G**ene****J****epa**

Test-time scaling in GeneJepa exploits a simple fact about the architecture. Reading depends on how many genes are shown, thinking does not. The encoder first reads an unordered set of genes into a fixed latent array using cross-attention, then a latent transformer reasons over that array at a cost that is independent of input size. This separation allows for exchanging compute for accuracy at inference without retraining the model or changing the encoder.

We partition each cell’s genes into a sequence of informative chunks, ordered by expression variability, and apply the same cross-attention block repeatedly. After the first chunk, the latents capture a coarse cellular state. Each additional chunk refines that state. The latent stack and the predictor run once at the end, exactly as in the standard forward pass. Because only the read stage changes, the extra work grows slowly while the dominant think stage stays fixed.

**Table 1:** GeneJepa improves directionality metrics relative to common baselines on the NeurIPS 2023 PBMC perturbation task. We report absolute GeneJepa performance and the improvement Δ over each comparator for cosine, Pearson, and Spearman.

| Method | Cosine |  | Pearson |  | Spearman |  |
| --- | --- | --- | --- | --- | --- | --- |
| | Value | $\Delta$ | Value | $\Delta$ | Value | $\Delta$ |
| GENEJEPa | **0.3698** | — | **0.3509** | — | **0.3431** | — |
| Mean outcome | 0.2264 | +0.1434 | 0.2198 | +0.1311 | 0.2117 | +0.1314 |
| Mean across compounds | 0.2628 | +0.1070 | 0.2594 | +0.0915 | 0.2425 | +0.1006 |
| Mean across cell types | 0.3017 | +0.0681 | 0.2972 | +0.0457 | 0.2806 | +0.0625 |
| Transformer ensemble | 0.2267 | +0.1431 | 0.2212 | +0.1297 | 0.2164 | +0.1267 |
| OP2 style baseline (JN-AP-OP2) | 0.3289 | +0.0409 | 0.3267 | +0.0242 | 0.3054 | +0.0377 |
| Random sample | 0.0547 | +0.3151 | 0.0524 | +0.2985 | 0.0562 | +0.2869 |
| Zeros | 0.0000 | +0.3698 | 0.0000 | +0.3509 | 0.0000 | +0.3431 |

In Figure 5, we see the expected scaling behavior on PBMC68k. Using a fixed probe trained on full embeddings, macro-F1 rises from about 0.02 with one read to about 0.35 with four reads, which matches the full model within 95% confidence intervals (Figure 5A). The embedding fidelity moves in lockstep, with cosine similarity approaching 1 as reads accumulate (Figure 5B). The wall clock per cell remains near 26ms and peak memory is essentially flat (Figure 5C). The result is a clean compute-to-quality frontier at inference that one can dial up when needed and leave unchanged when not. This capability follows directly from GeneJepa‘s Perceiver-style design. Inputs are sets rather than sequences, and the heavy latent computation is decoupled from input size. As a result, GeneJepa can refine its belief about a cell by rereading more genes while keeping the cost of thinking constant.

**Figure 5.**
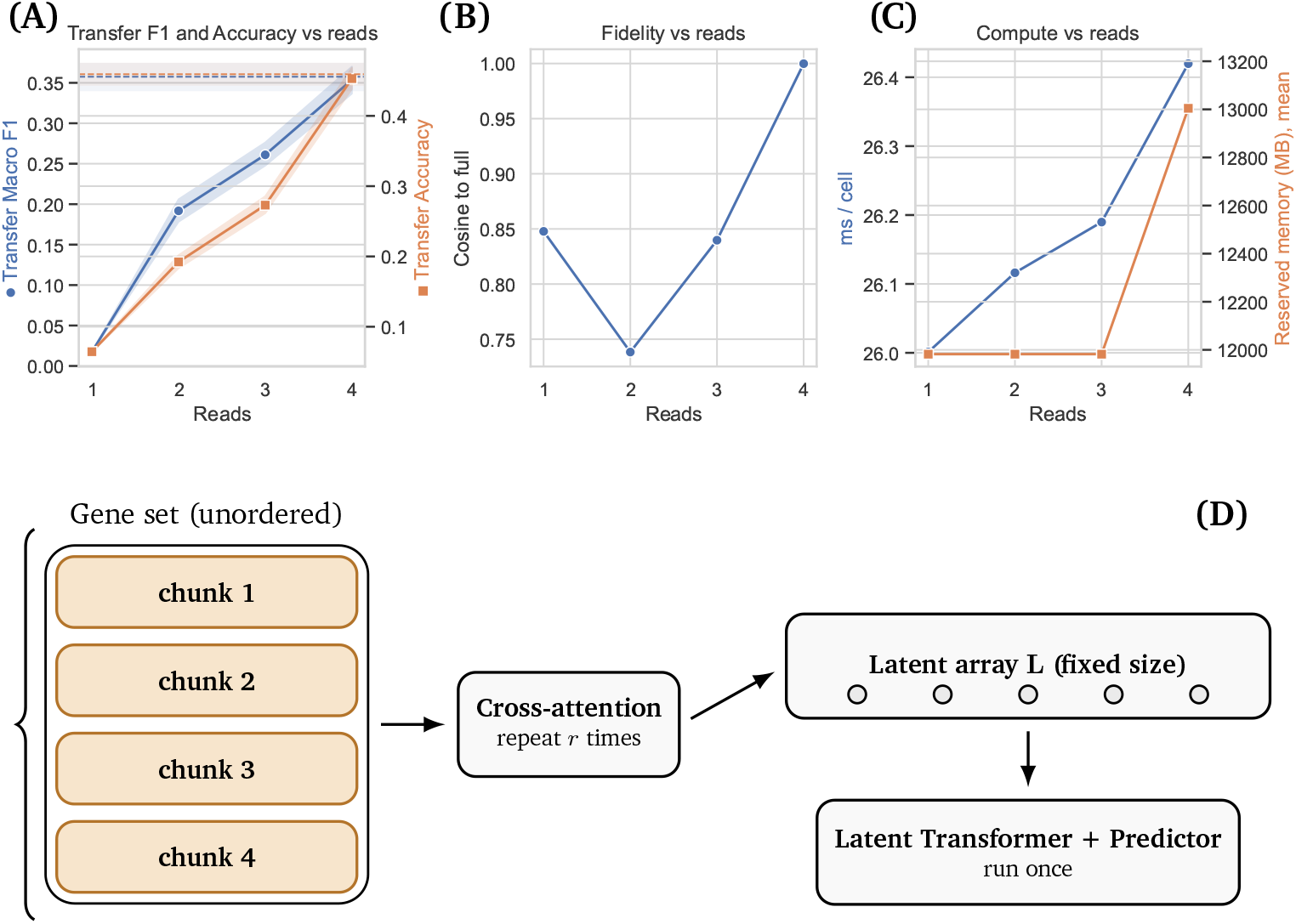
Multi-read test-time scaling in GeneJepa. Repeating the read over larger chunks improves fidelity and accuracy toward the full-input baseline while preserving a usable speed–quality trade-off via early exit. The latent computation is held fixed; only the read repeats. **(A)** Transfer Macro F1 and Accuracy vs. reads (shaded lines represent 95% confidence intervals). **(B)** Cosine to full embedding. **(C)** Inference time per cell and reserved memory vs. reads. A modest, one-time memory increase occurs only at the final read, which processes the largest input chunk. **(D)** Schematic of the iterative read pipeline. These trends show that GeneJepa is uniquely amenable to test-time scaling compared to existing single-cell foundation models, letting practitioners dial the number of reads to match time and memory budgets while recovering most of the full-input performance without retraining.

**Figure 6.**
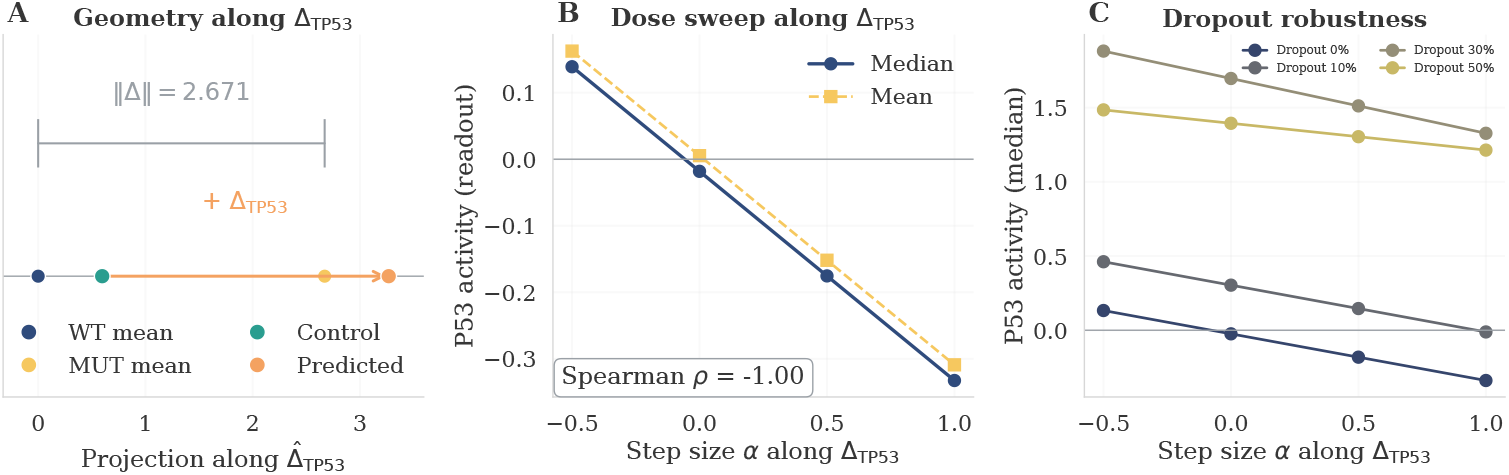
Zero-shot in-silico *TP53* knockout by GeneJepa. **(A)** Geometry along the learned perturbation direction Δ_TP53_: a WT control (green) moves to a predicted state (orange) along 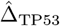; ∥Δ∥ = 2.671. WT and MUT means shown for reference. **(B)** Dose sweep: median (solid) and mean (dashed) P53-activity readout decrease monotonically with step size *α* (Spearman *ρ* = −1.00). **(C)** Robustness: monotonic trend persists under 0–50% input dropout, indicating a stable, interpretable perturbation effect. This shows that GeneJepa‘s latent space supports zero-shot in-silico gene knockout, without any perturbation-specific training.

### 4.5 Inverting the Predictive World Model to Simulate Zero-Shot Gene Knockout

Here we show that GeneJepa‘s embedding space is structured well enough that a gene knockout can be simulated by a single vector operation. To do this, we learn one direction per gene that points from wild-type to mutant-like states, then add that direction to any wild-type cell’s embedding to obtain a predicted post-knockout state. No perturbation pairs or training are required.

#### Creating a library of pertubation vectors

Let *f*_*θ*_ be the frozen GeneJepa encoder. For a gene *g*, we form two sets of cells from the large, heterogeneous Tahoe-100M corpus: a wild-type-like set *S*_WT_(*g*) and a mutant-like set *S*_MUT_(*g*). When cell-line metadata are available, we use them; otherwise we adopt a conservative proxy (quantiles of log-expression or pathway activity, stratified within cell line to reduce confounding). We then define the *knockout direction*

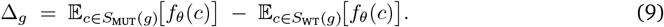

This is computed once per gene and reused. For *g* = `TP53`, the resulting direction has non-trivial length (∥Δ__TP53__∥ = 2.67) while the WT and MUT means remain close in angle (cos(*µ*_WT_, *µ*_MUT_) = 0.994), indicating a *small but coherent* displacement in the learned space.

#### Applying the knockout to a new cell

Given a wild-type control cell with embedding **z**_ctrl_, we simulate a knockout by

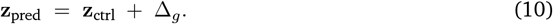

To quantify movement along the intended axis, we project onto the unit direction 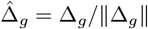 after centering at the WT mean *µ*_WT_:

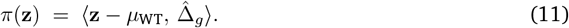

In our TP53 experiment, a representative control moves from *π*(**z**_ctrl_) = 0.60 to *π*(**z**_pred_) = 3.27 (gain = +2.67), that is, a clear shift toward the mutant mean along the *targeted* axis, not a generic drift.

#### Reading out biological consequences

We do not decode to counts. Instead we trained a single linear readout *s*(*z*) that maps embeddings to a P53 pathway activity score computed from the MSigDB HALLMARK_P53_PATHWAY gene set. The readout is a ridge regression fit once on embeddings and their pathway scores and then reused for all evaluations. It does not use perturbation pairs.

We compared a control embedding **z**_ctrl_ with the same embedding after adding the learned direction Δ_*g*_, denoted **z**_pred_. For `TP53`, **z**_pred_ lies closer to the proxy mutant mean than to the proxy wild type mean, with cos(**z**_pred_, *µ*_MUT_) = 0.994 and cos(**z**_pred_, *µ*_WT_) = 0.986, whereas the original control shows the opposite relation. To test for a graded effect, we traced the one dimensional path **z**(*α*) = **z**_ctrl_+*α* Δ_*g*_ and scored each point with the fixed readout. The score changed monotonically with *α*. Median values at *α ∈* {−0.5, 0, 0.5, 1} were 0.139, −0.018, −0.175, and −0.333, giving Spearman *ρ* = −1.0. To assess whether this trend is stable to noise, we randomly zeroed a fraction of the embedding coordinates and repeated the sweep. The trend persisted over rates from 0 to 0.5; absolute levels shifted as expected, but the slope remained stable. These observations show that adding the single vector Δ_*g*_ moves cells in the learned space toward a proxy mutant manifold and produces a consistent change in an external pathway readout that is trained once and reused for all cells.

Next, we complement the metadata-based construction of Δ_*g*_ with a label-free, zero-shot check that uses the same encoder *f*_*θ*_ and the same operations in Equation 9, Equation 10, and Equation 11. The only change is how Δ_*g*_ is estimated and how we evaluate it. On a build split, we remove gene *g* from each cell’s input, embed both the original cell and its ablated version, compute the per cell shift **z**^(0)^ − **z**, and define Δ_*g*_ as the average of these shifts. This uses no wild type or mutant labels, no proxies, and no perturbation pairs. On a separate held out split, we add this Δ_*g*_ to each unablated embedding and compare the result to that cell’s ablated embedding.

Table 2 reports the median percent reduction in the distance to the ablated target with a 95% bootstrap interval, the alignment cosine between the predicted shift and the true ablation shift, and the sample sizes. This demonstrates that the encoder supports a genuine zero-shot knockout, since a single learned direction reliably moves held out cells toward their ablated targets without labels or retraining.

**Table 2:** Zero-shot TP53 knockout in the GeneJepa latent space. On a held-out set of 20,000 cells, adding the zero-shot direction reduces the distance to the ablated embedding for a typical cell; we report effect sizes with uncertainty.

| Parameter | Value |
| --- | --- |
| Gene | TP53 |
| Build size | 30,000 cells |
| Evaluation size | 20,000 cells |
| Directional alignment (cosine) | 0.691 |
| Distance shrinkage (latent units, mean) | 0.0248 |
| Distance shrinkage (latent units, median) | 0.0358 |
| Percent shrinkage (median) | 27.17% ± 0.27% (95% CI) |
| Wilcoxon one-sided $p$ (median > 0) | $< 10^{-6}$ |

**Table 3:** GeneJepa Model Configuration.

| Parameter | Value |
| --- | --- |
| Embedding Dimension ($d$) | 768 |
| Number of Latents ($L$) | 512 |
| Number of Latent Blocks ($D$) | 24 |
| Number of Attention Heads ($h$) | 12 |
| Cross-Attention Chunk Size | 32 |
| Gene Vocabulary Size | 58,994 |
| Mask Ratio | 0.45 |
| Number of Target Blocks | 1 |
| Min. Context Genes | 512 |
| Min. Target Genes per Block | 16 |
| EMA Start Decay ($\beta_{\text{start}}$) | 0.992 |
| EMA End Decay ($\beta_{\text{end}}$) | 0.9995 |
| EMA Warmup Steps | 2000 |
| Predictor Depth | 3 |
| Predictor Expansion Factor | 4 |
| Tokenizer Identity/Value Split Ratio | 0.5 |
| Tokenizer Fourier Frequencies ($N_f$) | 64 |
| Fourier Min Frequency | 0.1 |
| Fourier Max Frequency | 100.0 |
| Fourier Frequency Scale | 1.0 |

**Table 4:** GeneJepa Training Configuration.

| Parameter | Value |
| --- | --- |
| Similarity Loss Coefficient ($\lambda_{\text{sim}}$) | 1.0 |
| Variance Loss Coefficient ($\lambda_{\text{var}}$) | 25.0 |
| Covariance Loss Coefficient ($\lambda_{\text{cov}}$) | 1.0 |
| Learning Rate | $1 \times 10^{-4}$ |
| Weight Decay | $2 \times 10^{-4}$ |
| Warmup Ratio | 0.05 |
| Max Epochs | 50 |
| Precision | bf16-mixed |
| Accumulate Gradient Batches | 2 |
| Gradient Clipping Value | 1.0 |
| Adam $\beta_1$ | 0.9 |
| Adam $\beta_2$ | 0.98 |

**Table 5:** Data and Experiment Configuration.

| Parameter | Value |
| --- | --- |
| Batch Size per Device | 92 |
| Number of Dataloader Workers | 8 |
| Effective Batch Size | $92 \times N_{\text{devices}} \times 2$ |
| Random Seed | 42 |

## 5 Conclusion

### Summary

GeneJepa introduces a new paradigm for genomics by learning a predictive world model of the transcriptome. We shift from signal-level reconstruction to abstract prediction in a latent space, forcing the model to learn the underlying rules of gene regulation rather than superficial data statistics. Our architecture, which combines a scalable Perceiver encoder with a continuous-value tokenizer, creates powerful representations from massive, unlabeled single-cell data. We demonstrate that these rep-representations are remarkably versatile; yielding state-of-the-art performance on cell-type identification across diverse tissues and accurately predicting cellular responses to unseen chemical perturbations. In contrast to prior single-cell foundation models, we show that our model’s learned structure can be inverted to perform zero-shot *in-silico* gene knockouts. By manipulating embeddings with pre-computed perturbation vectors, we can simulate transcriptome-wide effects of unseen genetic interventions, indicating that GeneJepa supports mechanistic inference in cell biology.

### Limitations

This work has three key limitations: (1) pretraining is dominated by cancer cell lines (Tahoe-100M), which may bias representations and limit transfer to primary tissues and non-cancer contexts; (2) the Jepa objective omits explicit batch-correction or domain-invariance terms, leaving robustness to lab effects to emerge implicitly from scale and inductive bias; and (3) our knockout analyses are evaluated in latent space only. Wet-lab studies are *sine qua non* for establishing biological realism.

### Outlook

A natural next step is to expand the scope of our world model by training on a more comprehensive gene set, moving beyond the 58,994-gene vocabulary of the current model to include the entire pan-genome of coding and non-coding elements. This would create a more complete and accurate simulation space. The true frontier, however, lies in creating multi-omic foundation models. The modality-agnostic nature of the Perceiver architecture is ideally suited for a multiomic GeneJepa, which could be trained to jointly predict the latent state of the transcriptome, epigenome, and proteome from one another. Concretely, one can pretrain a tri-omic GeneJepa that symmetrically predicts RNA latents from ATAC and surface-protein latents, and the reverse, through a shared Perceiver bottleneck, using modality dropout and a cycle-consistency penalty to leverage both paired and unpaired cells. Such a model would learn a unified, holistic representation of the cell, potentially uncovering the cross-modality regulatory logic that governs cellular function and disease. We believe that building and inverting these predictive world models represents the next logical step in computational biology, moving beyond statistical description and toward mechanistic and predictive understanding.

## Data availability and licensing

Tahoe-100M (Hugging Face: tahoebio/Tahoe-100M) is released under *CC0 1*.*0* (public-domain dedication). sci-Plex (v3) data are publicly available from NCBI GEO under Series accession GSE139944; GEO is an open-access archive and reuse follows GEO policies with citation of the originating study. HLCA data are available via the Human Cell Atlas Data Portal; downloaded and exported data are licensed under *CC BY 4*.*0* per the HCA Data Release/Data Use policies. PBMC3k (10x Genomics demonstration dataset) is distributed under *CC BY 4*.*0*; attribution is required.

## A Appendix

### A.1 Hyperparameter Details

This section provides the detailed hyperparameters used for the main GeneJepa model training and configuration.

#### A.2 Cross-Attention Implementation: Online Softmax (float32 accumulators)

##### Algorithm 1

**Stable Chunked Cross-Attention via Online Softmax (float32 accumulators)**

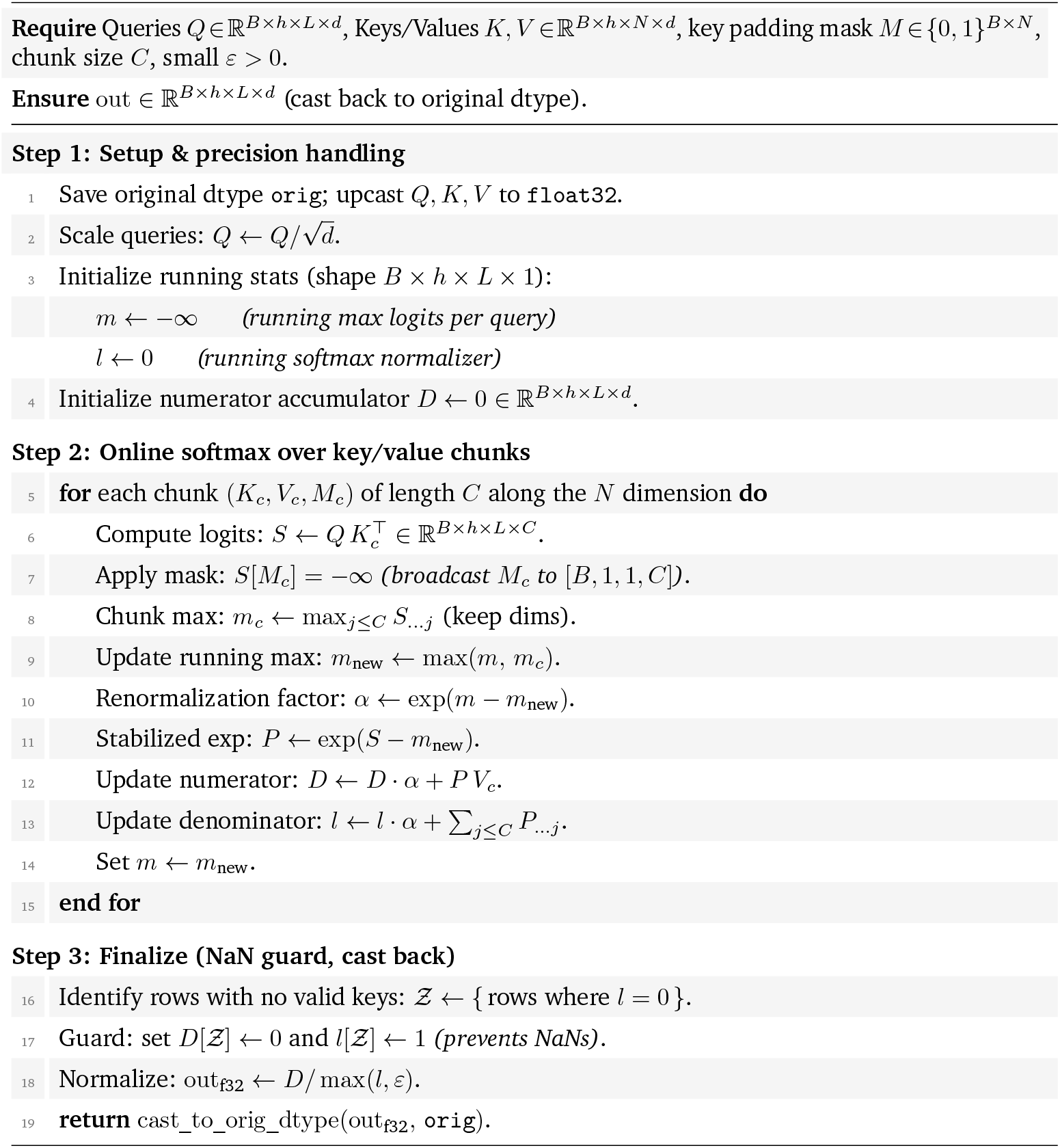

## Notes

https://github.com/BiostateAI/GeneJEPA

### Competing Interest Statement

EL, TM, VA, EM, OL, AG and TK are employees of Biostate AI. Additionally, TM, VA, OL, AG and TK are shareholders.

